# Novel *in vivo* TDP-43 stress reporter models to accelerate drug development in ALS

**DOI:** 10.1101/2023.12.18.572130

**Authors:** Febe Ferro, C. Roland Wolf, Chris Henstridge, Francisco Inesta-Vaquera

## Abstract

The development of therapies to combat neurodegenerative diseases is widely recognised as a research priority, with conditions like Alzheimer’s, Amyotrophic lateral sclerosis (ALS) and Parkinson’s set to place an ever-heavier burden on healthcare systems in the near future. Despite recent advances in understanding their molecular basis, there is a lack of suitable early biomarkers to test selected compounds and accelerate their translation to clinical trials. We have investigated the utility of *in vivo* reporters of cytoprotective pathways (e.g. NRF2, p53) as surrogate early biomarkers of the ALS degenerative disease progression. We hypothesized that cellular stress observed in a model of ALS may precede overt cellular damage and could activate our cytoprotective pathway reporters. To test this hypothesis, we generated novel ALS-reporter mice by crossing the hTDP-43tg model into our oxidative stress/inflammation (Hmox1; NRF2 pathway) and DNA damage (p21; p53 pathway) stress reporter models. Histological analysis of reporter expression in a homozygous hTDP-43tg background demonstrated a time-dependent and tissue-specific activation of the reporters in tissues directly associated with ALS. The activation occurs in Purkinje neurons and other parvalbumin-positive (PV+) cells within the cerebellum of mice, before moderate clinical signs are observed. In addition, reporter expression in hTDP-43tg hom peripheral tissues was not observed at the tested mouse ages (15 and 17 days postnatally). Further work is warranted to determine the specific mechanisms by which TDP-43 accumulation leads to reporter activation and whether therapeutic intervention modulates reporters’ expression. Our current studies suggest that these reporters may represent a powerful approach to accelerate preclinical studies targeting TDP-43 pathologies. We anticipate the reporter strategy could be of great value in developing treatments for a range of degenerative disorders.

## Introduction

Age-related diseases account for a high proportion of the total global burden of disease. Degenerative diseases of old age, including cardiovascular disease, stroke, type II diabetes, and chronic respiratory conditions, have emerged as the major causes of death worldwide. Of special concern are neurodegenerative conditions including Alzheimer’s (AD), Parkinson’s disease and other dementias, which have been described as the greatest unmet need facing modern medicine (1,2). Developing new therapies to slow, halt, or reverse these pathologies is regarded as a key element in addressing this global health priority. Unfortunately, no prognostic biomarkers of degeneration are available to predict whether a candidate disease-modifying therapy is likely to improve clinical outcome (3). As a consequence, there is an urgent need for prognostic biomarkers of degeneration that can improve the ability of pre-clinical research to correctly predict future clinical benefit (4).

One major impediment in developing new treatments of these diseases is the lack of robust models in which identify and prioritize therapeutic approaches to be tested and then progressed to clinical trials in a timely manner. The development of early biomarkers prior to the onset of clinical symptoms would at least in part address this issue (5).

ALS is a progressive neurodegenerative disease, characterized by motor neuron loss, which results in paralysis and death 3-5 years after disease onset (6). At the molecular level, most sporadic and familial ALS cases are characterized neuropathologically by cellular aggregates of Transactive response DNA-binding Protein 43 kDa (TDP-43) (7). Furthermore, TDP-43 aggregates are observed in approximately 50% of Frontotemporal dementia (FTD), 30% AD and 20% Parkinson’s disease cases, so understanding its toxic mechanism(s) is essential. The mechanism of TDP-43 induced cell death is associated with the induction of cellular stress, however the actual stress responses involved (e.g. inflammation, oxidative stress, DNA damage), remain unclear (8). There is therefore a need to characterise these pathways to aid the development of new treatments for this intractable disease (9).

The mechanisms of TDP-43 aggregation and its associated toxicity (inflammation, oxidative stress and DNA damage) in ALS/FTD are still poorly understood (7,9,10). To improve our understanding of TDP-43 toxicity, researchers have generated several TPD-43 murine models that mimic human ALS/FTD pathologies. One of these models, the hTDP-43tg, overexpresses human wild-type TDP-43 under the control of the neuronal murine Thy-1 promoter that drives transgene expression in virtually all neurons of the central nervous system from 1 week after birth (11,12). Homozygous hTDP-43tg mice express a 4 to 5-fold induction of hTDP-43 compared with murine TDP-43. Whereas mice hemizygous for the TDP-43 transgene (TDP-43^het^) are viable, fertile, and grossly normal, mice that have two copies of this transgene (TDP-43^hom^) display profound motor dysfunction, resulting in an inability to walk around P21 and death around P24. This model provides an opportunity to study TDP-43-driven pathology and cell stress in a short timeframe.

We have developed a portfolio of reporter mouse models in which the activation of cellular cytoprotective pathways, including inflammation/oxidative stress (NRF2-Hmox1 reporter) and senescence/DNA damage (p53-p21 reporter), can be monitored at single-cell resolution across all organs, tissues, and cell types (13–15). Moreover, the measurements can also be performed non-invasively and in real-time. In each model, a short viral DNA sequence, known as a 2A sequence, is exploited to provide multiple reporters expressed (separately) from a single gene promoter. For example, when Hmox1 reporter mice were exposed to inducers of oxidative stress (e.g. paracetamol) or inflammatory mediators (e.g. LPS), a tissue- and cell-specific induction of ß-galactosidase reporter activity was observed. Similarly, when the p21 reporter mice were exposed to DNA damage inducing agents (e.g. gamma-irradiation or cisplatin) an increase of p21 expression was detected at a high level of fidelity and resolution in target tissues. We therefore wanted to explore whether cellular stress resulting from accumulation of hTDP-43, including DNA damage and inflammation, would activate the expression of these reporters in neurons.

In this short communication we report the generation of a hTDP-43 reporter system that could be applied to test the efficacy of novel therapeutic interventions for motor neuron disease. These mice will be made available to the research community.

## Methods

### Animals

All animals used in this study were bred and maintained in the CIR Resource Unit, School of Life Sciences, University of Dundee. hTDP-43 animals were purchased from Jackson Lab (B6;SJL-Tg(Thy1-TARDBP)4Singh/J; strain number 012836) and they were described before (11). hTDP-43_p21 and hTDP-43_Hmox1 reporter mice were generated by crossing hTDP-43 heterozygous (hTDP43^het^) mice into heterozygous Hmox1 (HOTT^het^) or p21 (p21^het^) reporter mice (13). Mice were housed in open-top cages in temperature-controlled rooms at 21°C, with 45-65% relative humidity and 12h/12h light/dark cycle. Mice had ad libitum access to food (R&M No.1 for stock females; R&M No. 3 for mating females; Special Diet Services, Essex, UK); and water. Animals were regularly subjected to health and welfare monitoring as standard (twice daily). Environmental enrichment was provided for all animals. All animal work described was approved by the Welfare and Ethical use of Animals Committee of the University of Dundee. No regulated procedures were conducted in these animals.

The severity of hTDP-43 mice phenotype was assessed according to the established scoring system. Briefly, a score of 0 correspond to mice that when suspended by the tail and both hindlimbs were consistently splayed outward, away from the abdomen. At this score system, mouse moves normally with body weight supported on all limbs; a score of 1 was achieved when mice suspended by the tail retracted one hindlimb toward the abdomen for more than 50% of the time suspended. Mice at this score had a mild tremor or a limp while walking; a score of 2 was given to those mice that when suspended by the tail, both limbs are partially retracted towards the body for 50% of the time suspended. Mice at this score show severe tremor and/or limp, or the feet point away from the body during locomotion (“duck feet”); finally, a score of 3 is reached when mice are suspended by the tail, both limbs are fully retracted for more than 50% of the time suspended and/or the mouse has difficulty moving forward and drags its abdomen along the ground. Throughout our studies, no animals progressed to score 2.

### Genotyping

Genotyping for the p21 and HOTT strains was performed as described before (16), and according to instructions in the case of hTDP-43 mice. The primers used were as follows: Hmox1 reporter: HO1-KI Fwd, 5’-GCTGTATTACCTTTGGAGCAGG-3’; HO-1-KI Rvr, 5’-CCAAAGAGGTAGCTAATTCTATCAGG-3’; p21 reporter: p21-KI Fwd, 5’-GCTACTTGTGCTGTTTGCACC-3’; p21-KI Rvr, 5’-TCAAGGCTTTAGGTTCAAGTACC-3’); hTDP-43 common primer: 5’-TGAAATCCGGGTGGTATTGG-3’; hTDP-43 (wild type allele): 5’-GGTGAGTTTAACCTTCAAGGGCT-3’; hTDP-43 (transgene): 5’-AGCTTGCTAGCGGATCCAGAC-3’.

### Tissue harvesting and processing for cryo-sectioning

Mice were perfused with PBS followed by 4% paraformaldehyde in 0.1 M PBS pH 7.2-7.4 (flow rate 5mL/min). Brains were harvested, halved, and post-fixed in 4%PFA overnight for IHF and for 3 hours for LacZ staining. Brains were sectioned using a vibratome (Leica VT1000S). Superglue (LocTite SuperAttak Gel) was used to attach the brain to the cutting surface vertically, so the cerebral cortex is facing up. Tissues are immersed in a PBS bath. Blades Gillette “Wilkinson Sword” were used. Sections were cut at 40 μm thickness and saved in a 24-well tissue-culture dish filled with 1X PBS, 0.5% sodium azide (5 sections/well). Sections were picked up with a paintbrush. One section per well was mounted on slides for IHC. Tissues preserved for β-galactosidase staining were rapidly harvested postmortem and processed by immersion fixation in 4% paraformaldehyde (PFA) (brain, small intestine) for 2h, 3% neutral-buffered formalin (NBF) (liver) for 3h or Mirsky’s fixative (rest of tissues) for 24h and subsequently cryoprotected for 24h in 30% (w/v) sucrose in phosphate-buffered saline (PBS) at 4°C. Organs were embedded in Shandon M-1 Embedding Matrix in a dry ice-isopentane bath. Sectioning was performed on an OFT5000 cryostat (Bright Instrument Co.). With the exception of lung (14μm) and brain (20μm) sections, all sections were cut at 10μm thickness.

### In situ β-galactosidase staining and histochemistry

Sections were rehydrated in PBS at room temperature for 15 minutes before being incubated overnight at 37°C in X-gal staining solution: PBS (pH 7.4) containing 2 mM MgCl2, 0.01% (w/v) sodium deoxycholate, 0.02% (v/v) Igepal-CA630, 5 mM potassium ferricyanide, 5 mM potassium ferrocyanide and 1 mg/ml 5-bromo-4-chloro-3-indolyl β-D-galactopyranoside. On the following day, slides were washed in phosphate buffer solution, counterstained in Nuclear FastRed (Vector Laboratories) for 4 min, washed twice in distilled water for 2 minutes and dehydrated through 70% and 95% ethanol (4.5 and 1 minute respectively) before being incubated in Histoclear (VWR) for 3 minutes, air-dried and mounted in DPX mountant (Sigma). Because the β-gal protein contains a nuclear localization signal it is localized specifically in cell nuclei (17).

### Tissue Immunofluorescence

Tissue slices were washed in 0.1M phosphate buffer before submerging in 10% sucrose for 15min followed by 30% sucrose at room temperature for 3hrs. Slices were wrapped in tin foil and permeabilization was performed by four freeze/thaw cycles above liquid nitrogen and finally placed in 0.1M phosphate buffer. After peroxidase blocking with 1% H_2_O_2_ slices were washed in 0.1M phosphate buffer and tris buffered saline solution and antigen blocking for 45 minutes at room temperature with 1% Human Serum Albumin. Primary antibodies were incubated overnight at 4°C before TBS washing and incubation with secondary antibody (1h room temperature). After phosphate buffer washing, sections were allowed to slightly dry and mounted with Vectashield. Primary antibodies used: TDP-43 (Abcam, ab133547 Rabbit monoclonal [EPR5811]); GFAP (Proteintech, 16825-1-AP Rabbit/IgG); Parvalbumin (Swant, PV27 rabbit).

### Image acquisition and quantitation

Images from immunolabelled tissues were acquired using a Delta Vision Elite High Resolution Microscope with a Leica 20X APO 1.4 lens. Convoluted Z-stacks images were acquired from multiple fields. Exposure and laser intensity was fixed before proceeding with fluorescence quantification to allow direct comparisons between sections. ImageJ (v2.0.0, NIH) analysis software was used for relative fluorescence intensity quantitation. Image analysis was performed as previously described (McCloy RA). Briefly, colour channels were split, and regions of interest (ROI) were selected. Measurement values for area, integrated density, mean/max/min grey value were recorded. A minimum of ten background ROIs were selected. Corrected total cell fluorescence (CTFC) was calculated = Integrated Density – (Area of selected cell x Mean fluorescence of background readings) (18). CTCF values were obtained by measuring the fluorescence intensity of TDP-43 in selected cells across three different fields of view in slides from individual experimental animal. The obtained values were then averaged for each ROI, and subsequently plotted as hTDP-43^wt^_HOTT^het^ vs. hTDP-43^hom^_HOTT^het^ and statistical analysis (1-sided unpaired Student *t* tests) was performed using GraphPad Prism 10.0.2.

## Results and Discussion

To establish whether the *in vivo* reporters of DNA damage (p21) or oxidative stress/inflammation (Hmox1) act as early biomarkers of TDP-43 induced pathologies, hTDP-43 mice were crossed into HOTT^hom^ as well as p21^hom^ reporter mice resulting in the TDP-43_HOTT or TDP-43_p21 lines. These mice were used to generate hTDP-43^hom^_HOTT and hTDP-43^hom^_p21 mice. The average litter sizes resulting from these crosses was ∼ 8 +/-2 pups and within normal mendelian ratios. For ethical reasons, mice were only kept until they were 17 days old. At this stage, hTDP-43^hom^ mice showed an expected phenotype of mild tremor and or a limp while walking (score 1). A complete list of collected tissues from different genotypes and animal age is provided (Table S1).

Histological analysis of PND17 TDP-43^hom^-HOTT^het^ reporter mice demonstrated a profound induction of reporter expression in a disease relevant tissue, the cerebellum (Figure 1) (19–21). The basal reporter expression and the increased signal due to hTDP-43 accumulation was similar in male and female so animals of both sexes were included in the reporter analysis of different age groups. We found a cell specific activation of the reporter in the cerebellum and typically in what appear to be PV+ cells, such as Purkinje (Purkinje layer) and basket (molecular layer) cells. Immunofluorescence staining of hTDP-43wt or hTDP-43hom cerebellum samples with parvalbumin (general GABAergic cell marker, including Purkinje and basket cells) and GFAP (astrocytes) markers confirmed the likely identity of these cells in the reporter-positive region (Figure S1A). In addition, quantitative analysis of hTDP-43 immunofluorescence staining demonstrated a 2-fold increased expression of the transgene in this area, consistent with previous expression analysis reports for this line (Figure S1B) (11).

**Figure 1.**
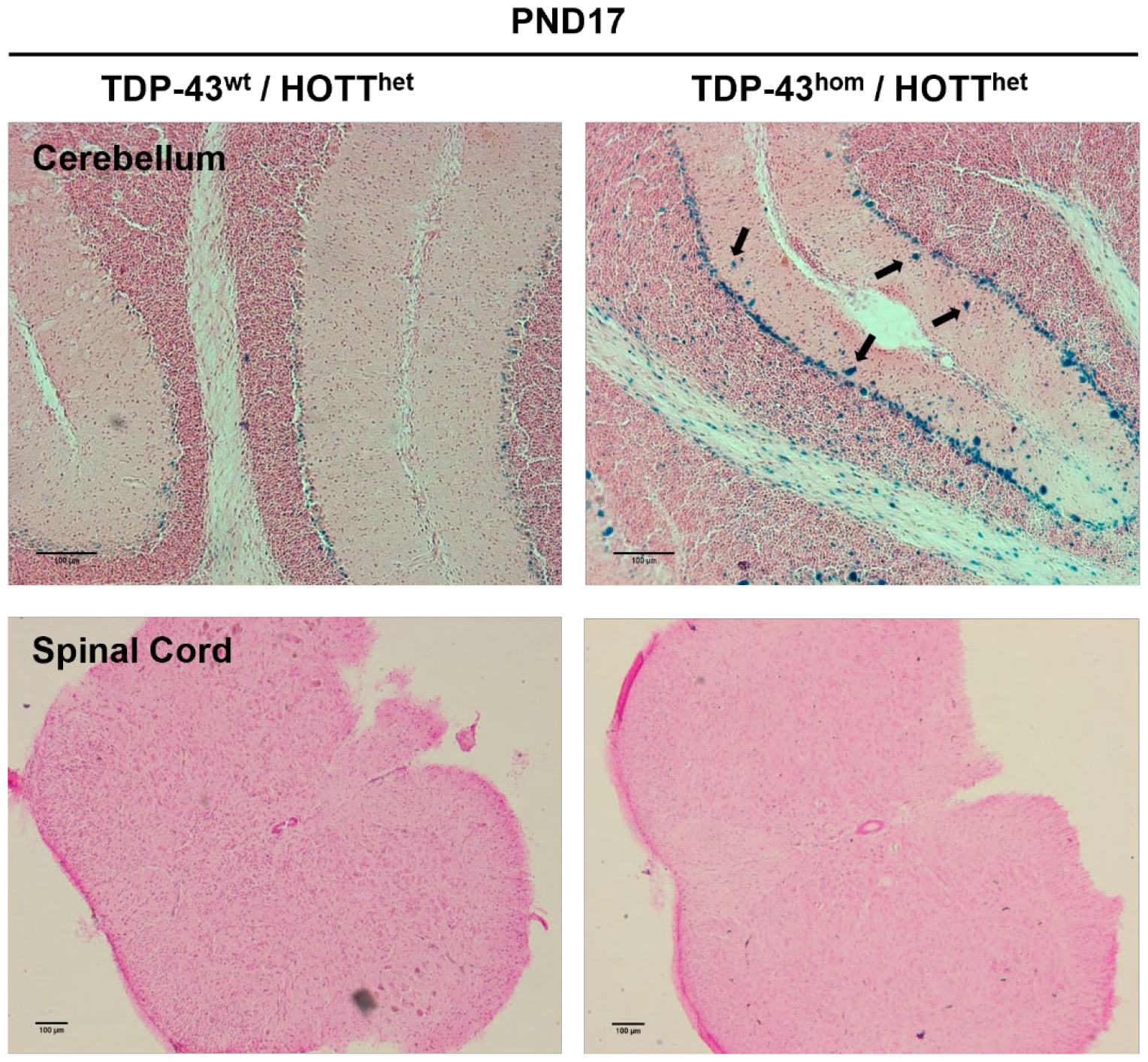
Demonstration of the HOTT reporter model utility to detect cellular stress associated with hTDP-43 accumulation. Representative images of LacZ staining in cerebellum (upper row) or lumar spinal cord (lower row) sections from triplicated mice of indicated genotypes. PND17, post-natal day 17. Black scale bar: 40 μm. Black arrows indicate positive cells for LacZ staining.

We extended our reporter expression analysis to other disease relevant areas of the central nervous system (CNS) and peripheral organs. These included the spinal cord, cortex and hippocampus (Figure 1 and Figure S2). Positive β-gal staining was observed in hTDP-43^wt^-HOTT^het^ mice as previously described in dentate gyrus and cortex regions (13). Differential expression of the reporter was not observed between the experimental groups in areas of spinal cord (Figure 1). However, the interpretation of these data was confounded by the fact that a variable reporter expression in this tissue was observed irrespective of the genotype. The significance of basal Hmox1 expression in these cells warrants further investigation.

The reporter activation in hTDP-43^hom^ mice occurred at a clinical score of 1, i.e. which precedes overt morbid phenotypes. Therefore, reporter activation provides a more refined endpoint to assess the effectiveness of therapeutic interventions aimed at reducing hTDP-43 accumulation (22). In addition, because changes in reporter expression were consistent between individual mice, the number of mice required to obtain statistical significance relative to current approaches is reduced. We would caveat that the β-gal reporter assay should be interpreted as a qualitative marker for pathway activation, which does not inform on the strength of the response. Indeed, when we co-stained hTDP-43^hom^_HOTT^het^ cerebellum sections with a β-gal antibody we didn’t observe a robust signal, suggesting that the expression of the β-gal reporter is below the limits of antibody-based detection. As a staining control, we observed positive β-gal in the nucleus of liver cells in samples previously generated in our lab (not shown).

To investigate whether reporter activation occurs at earlier time points, we analysed samples derived from PND15 pups in hTDP-43^hom^-HOTT^het^ and hTDP-43^hom^-p21^het^ mice (Figure 2 and S4). A mild but consistent activation of the reporter in the cerebellum of hTDP-43^hom^-HOTT^het^ mice was detected, similar of that observed at later stages. Unfortunately, the very high basal expression of the p21 reporter in the cerebellum precluded robust results with this mouse line. However, we observed distinctive positive (potentially basket) cells in the cerebellum of p21 reporter mice, similar to those observed in the HOTT line, suggesting this pathway is also activated in hTDP-43^hom^ mice (Figure 2). No differential reporter activation was observed in other areas of the brain (Figure S4).

**Figure 2.**
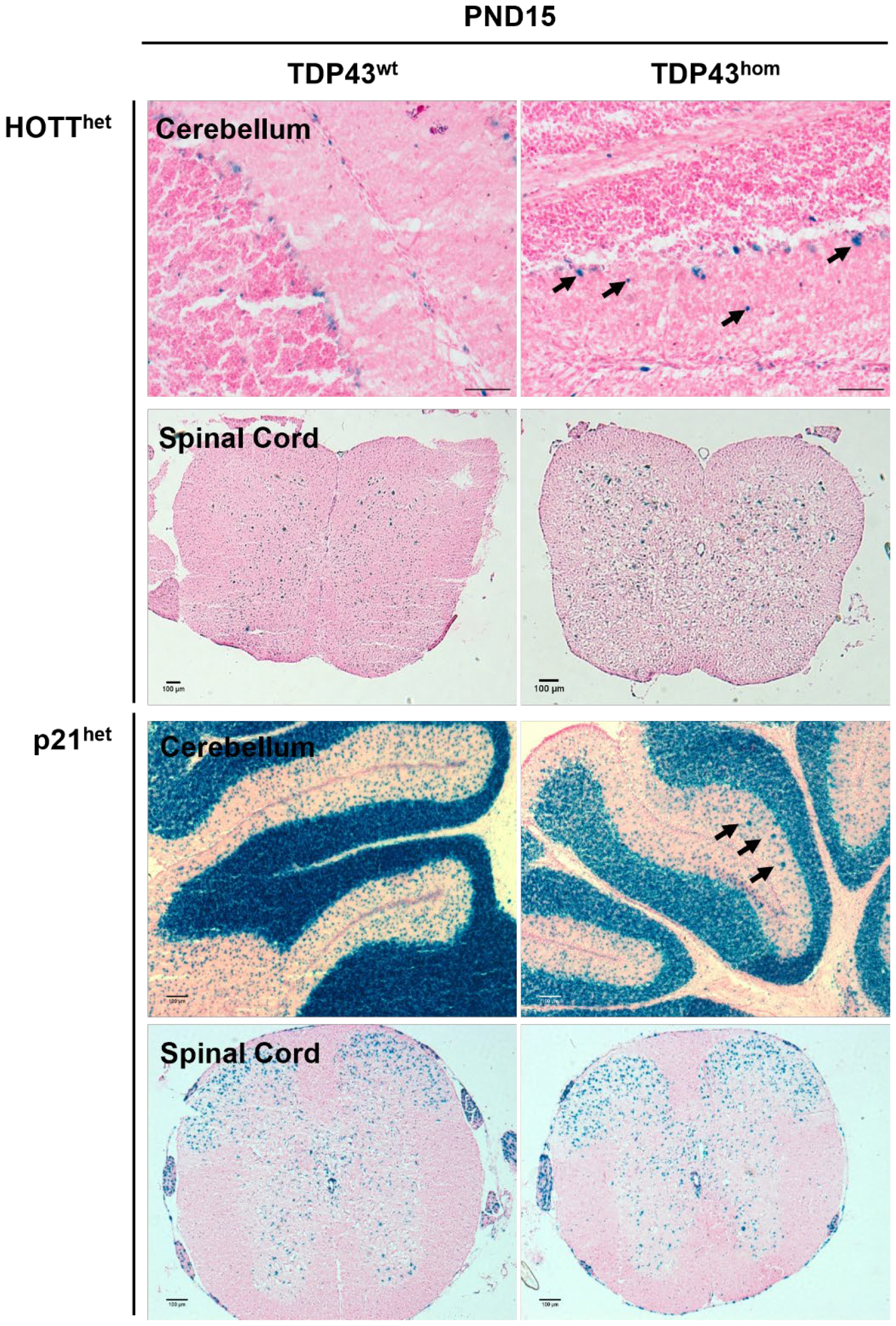
Early detection of hTDP-43 associated stress responses using the HOTT and p21 reporter models. Representative images of LacZ staining in cerebellum and spinal cords sections of Hmox1 (upper rows) or p21 (lower rows) reporters from triplicated mice of indicated genotypes. PND15, post-natal day 15. Black scale bar: 100 μm. Black arrows indicate positive cells for LacZ staining.

Remarkably, we have previously observed an enhanced sensitivity of PV+ cells in a novel TDP-43 mouse with a single point mutation inserted within the mouse TDP-43 gene (TDP-43^Q331K/Q331K^)(23). In this model, a reduction in the number of parvalbumin interneurons correlated with regional brain atrophy in mutant mice at both 5 and 20 months of age. Interestingly, this model also presents with early cerebellar atrophy, so together, we present evidence of a potential contribution of cerebellar PV+ cells to the pathogenesis of ALS. Further work is warranted to investigate whether the activation of the Hmox1 (oxidative stress/inflammation) or p21 (DNA damage) pathways are molecularly linked to the loss of PV cells in other TDP-43 mouse models and ALS patients.

We further investigated a potential hTDP-43 transgene dosage effect. We analysed hTDP-43het reporter expression in tissues of mice up to 40 weeks of age. No changes in β-gal staining were observed in any of the tissues studied, including in the cerebellum, spinal cord, cortex and hippocampus (Figure S4).

The data from our studies demonstrate the activation of specific TDP-43 dependent cell stress response pathways in a disease relevant cell population (20,21,24,25). It is important to note that the toxicity of severe phenotypes can lead to simultaneous activation of multiple stress pathways (toxicity burst), making it difficult to define the molecular hierarchy of these events (26). Our reporter data demonstrates the early induction of oxidative stress, inflammation (pathways leading to Hmox1) and DNA damage (p21 reporter) in TDP-43 pathologies. However, other cellular stress responses have been associated with the accumulation of hTDP-43, including integrated stress response (27), endoplasmic reticulum (ER) stress and the unfolded protein response (UPR) (28). Therefore, whether the pathways leading to Hmox1 or p21 expression are a cause or a consequence of TDP-43 related changes remains to be studied. Nevertheless, it will be important to identify the exact mechanism(s) linking hTDP-43 dependent and independent expression with the reporter activation in critical ALS related brain areas. Indeed, some therapies being developed to treat ALS are targeted at the NRF2 pathway, which is a major regulator of the Hmox1 gene (29).

In summary, this work demonstrates the potential use of our stress reporter approaches to accelerate finding new treatments for early stages of degenerative diseases prior to any perceptible deleterious phenotype. We will support the use of hTDP-43-HOTT mice as a new tool for interventional studies in TDP-43-dependent diseases. The accessibility of these models will accelerate testing molecules / approaches and their PK profiles before human trials.

## Acknowledgements

This research was funded by an Alzheimer’s Research UK (ARUK-PPG2020A-003) and MND Scotland (RC21-06) pilot grants awarded to FIV/CH.

We thank Prof. Tom Gillingwater and Dr. Helena Chaytow for discussions, genotyping, and animal scoring advice. We also thank Chris Henstridge lab members for their advice on tissue processing and analysis.

---

**Table S1.** Genotypes, age, and tissues collected during the project.

| Genotype | Age | Tissues Collected |
| --- | --- | --- |
| hTDP-43<sup>wt</sup>_HOTT<sup>het</sup> | 40 weeks | Brain, Spinal Cord,<br>Gastrocnemius,<br>Heart, Lungs,<br>Liver, Kidneys,<br>Spleen, Large<br>Intestine, Plasma |
|  | 24 weeks |  |
|  | PND17 |  |
|  | PND15 |  |
| hTDP-43<sup>het</sup>_HOTT<sup>het</sup> | 40 weeks |  |
|  | 24 weeks |  |
|  | 12 weeks |  |
|  | PND17 |  |
| hTDP-43<sup>hom</sup>_HOTT<sup>het</sup> | PND17 |  |
|  | PND15 |  |
| hTDP-43<sup>hom</sup>_HOTT<sup>hom</sup> | PND15 |  |
| hTDP-43<sup>wt</sup>_HOTT<sup>hom</sup> | PND17 |  |
| hTDP-43<sup>wt</sup>_p21<sup>het</sup> | 40 weeks |  |
|  | 24 weeks |  |
|  | PND15 |  |
| hTDP-43<sup>het</sup>_p21<sup>het</sup> | 40 weeks |  |
|  | 24 weeks |  |
|  | 12 weeks |  |
| hTDP-43<sup>hom</sup>_p21<sup>het</sup> | PND15 |  |
| hTDP-43<sup>hom</sup>_p21<sup>hom</sup> | PND15 |  |

**Figure S1.**
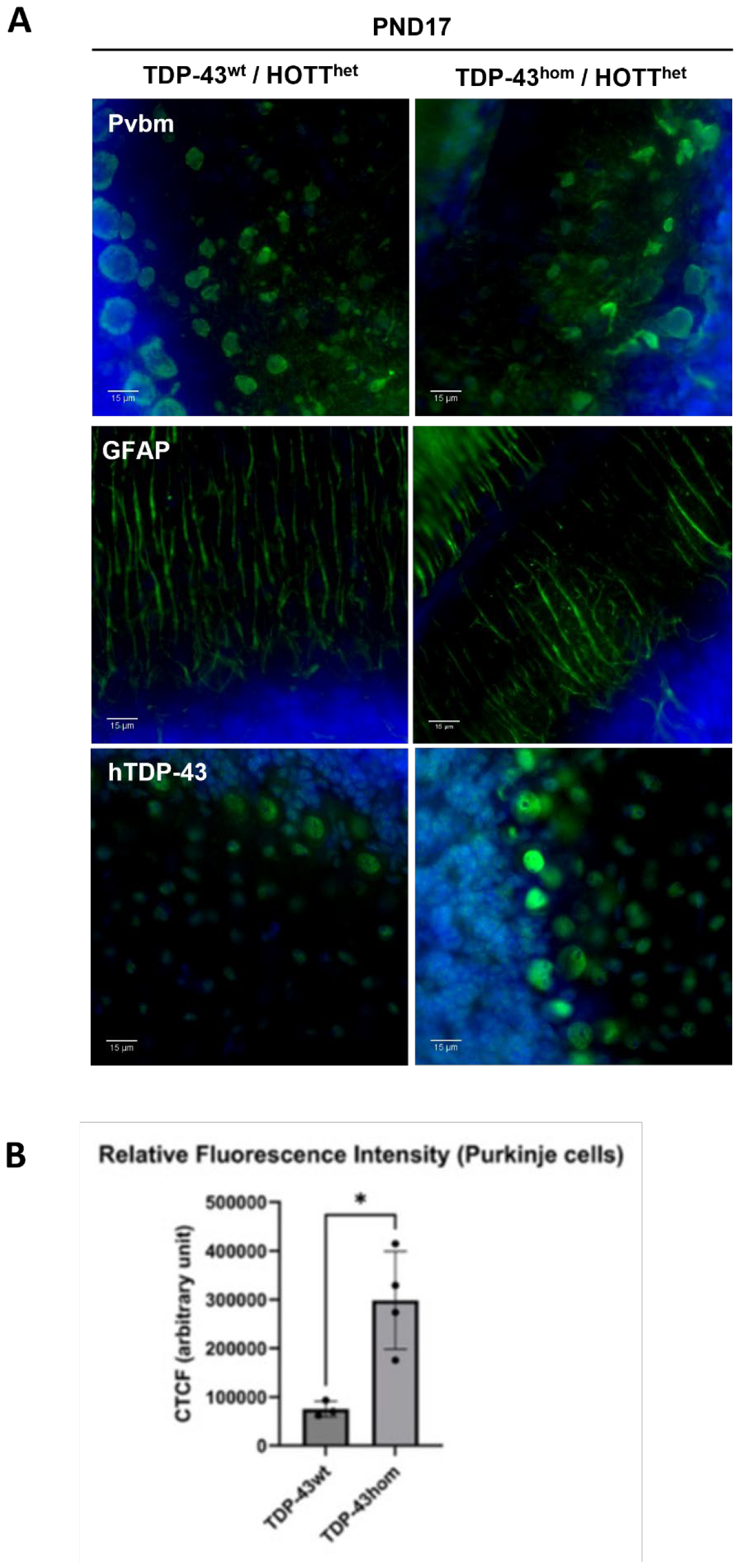
hTDP-43 accumulates in the cerebellum of homozygous mice. **A**. Staining with indicated antibodies in cerebellum samples from post-natal day 17 mice (PND17). Pvbm: parvalbumin; GFAP: glial fibrillary acidic protein; Green: primary antibodies. Blue: nuclear staining. White scale bar: 15 μm. **B**. Quantitation of relative hTDP-43 fluorescence intensity in Purkinje cells. * = p < 0.05.

**Figure S2.**
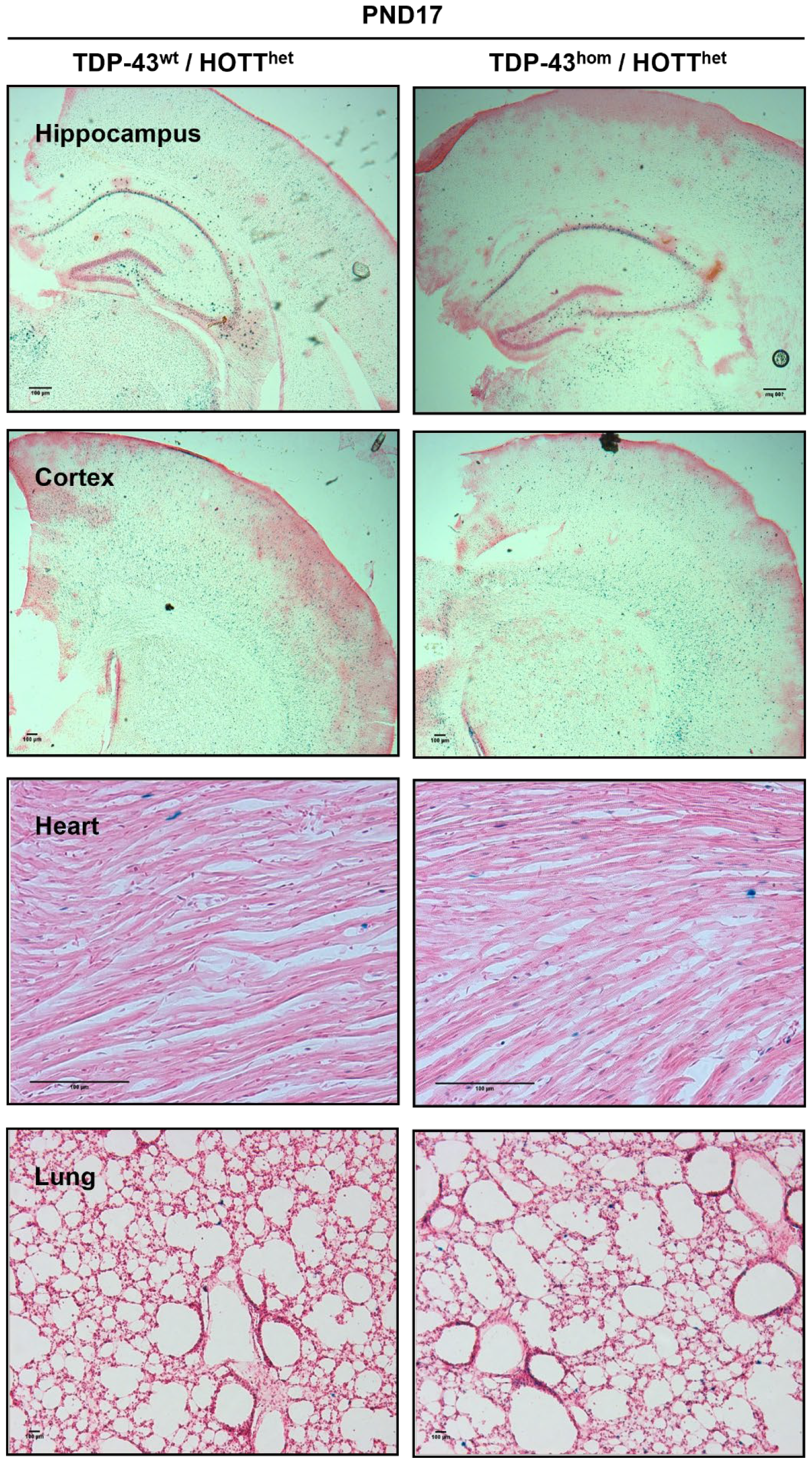
Absence of hTDP-43-related HOTT reporter expression in other areas of the brain or peripheral tissues at PND17. Representative images of LacZ staining at indicated brain areas, heart or lung tissues from triplicated mice of indicated genotypes. PND17, post-natal day 17. Black scale bar: 100 μm.

**Figure S3.**
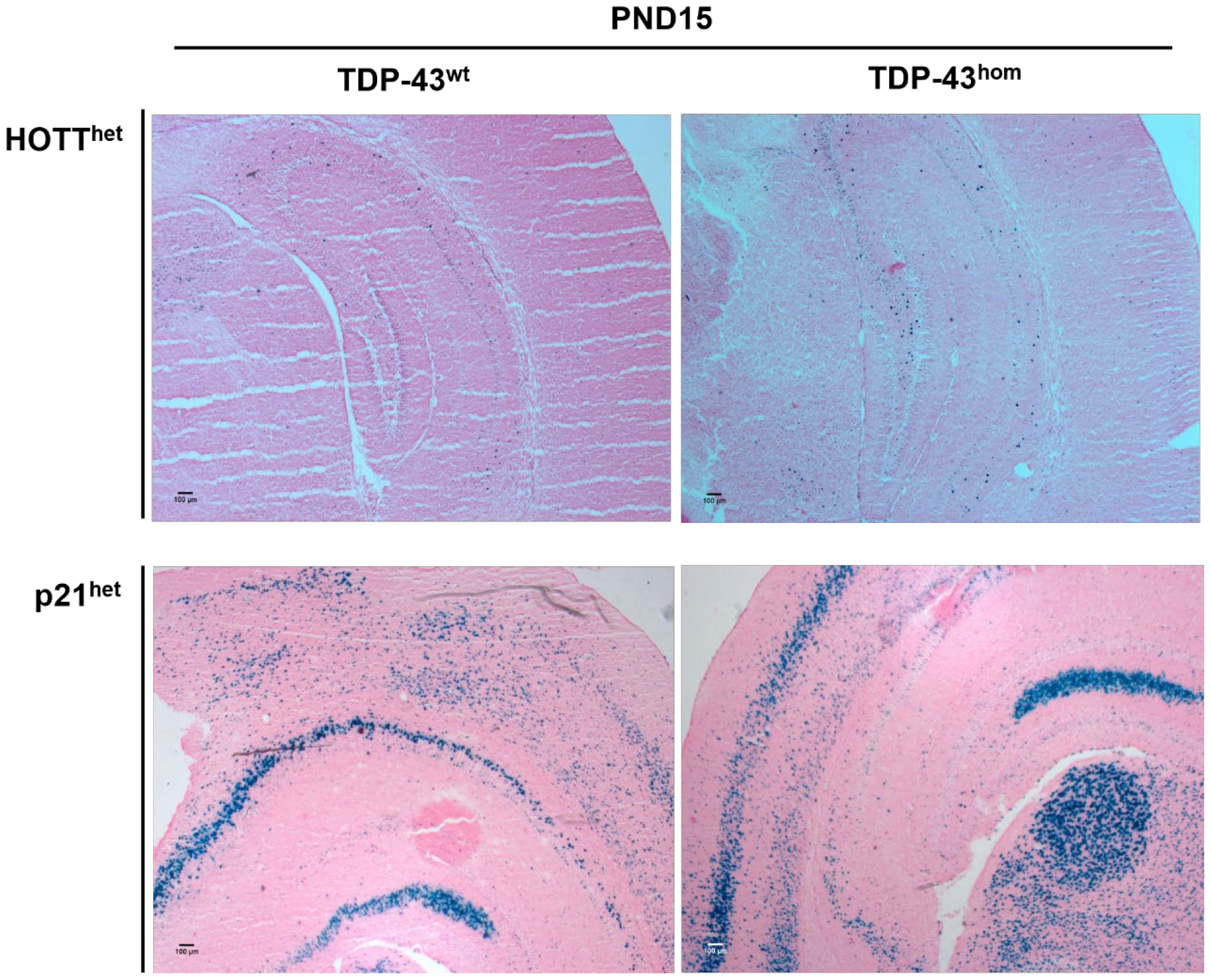
Absence of hTDP-43-related HOTT/p21 reporters’ expression in other areas of the CNS at PND15. Representative images of LacZ staining at cortex/hippocampus brain areas. PND15, post-natal day 15. Black scale bar: 100 μm.

**Figure S4.**
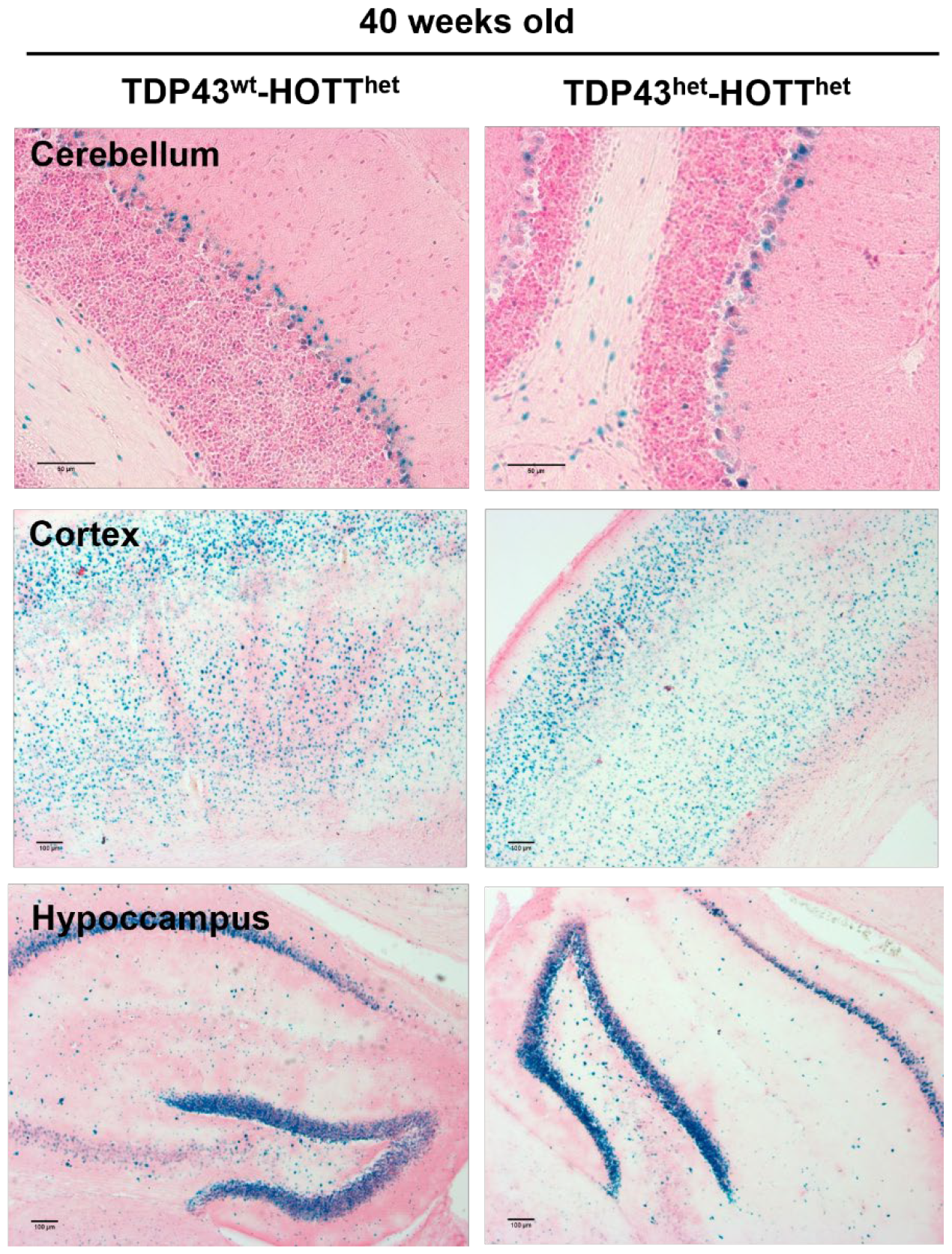
Absence of Hmox1 reporter expression changes in hTDP-43 heterozygous mice. Representative images of LacZ staining in brain sections from triplicated mice of indicated genotypes. Black scale bar: 50 μm (cerebellum sections); 100 μm (cortex and hippocampus). Black arrows indicate positive cells for LacZ staining.

## Bibliography

1. Feigin VL, Vos T, Nichols E, Owolabi MO, Carroll WM, Dichgans M, et al. The global burden of neurological disorders: translating evidence into policy. Lancet Neurol [Internet]. 2020 Mar 1 [cited 2023 Oct 24];19(3):255–65. Available from: http://www.thelancet.com/article/S1474442219304119/fulltext

2. Feigin V, Vos T. Global burden of neurological disorders: from Global burden of disease estimates to actions. Neuroepidemiology. 2019;52:1–2.

3. Hansson O. Biomarkers for neurodegenerative diseases. Nature Medicine 2021 27:6 [Internet]. 2021 Jun 3 [cited 2023 Oct 24];27(6):954–63. Available from: https://www.nature.com/articles/s41591-021-01382-x

4. Petrov D, Mansfield C, Moussy A, Hermine O. ALS clinical trials review: 20 years of failure. Are we any closer to registering a new treatment? Front Aging Neurosci. 2017 Mar 22;9(MAR):68.

5. Tan RH, Ke YD, Ittner LM, Halliday GM. ALS/FTLD: experimental models and reality. Acta Neuropathol. 2017 Feb 1;133(2):177–96.

6. Masrori P, Van Damme P. Amyotrophic lateral sclerosis: a clinical review. Eur J Neurol. 2020 Oct 1;27(10):1918–29.

7. Neumann M, Sampathu DM, Kwong LK, Truax AC, Micsenyi MC, Chou TT, et al. Ubiquitinated TDP-43 in frontotemporal lobar degeneration and amyotrophic lateral sclerosis. Science (1979). 2006 Oct 6;314(5796):130–3.

8. Mackenzie IRA, Neumann M. Molecular neuropathology of frontotemporal dementia: insights into disease mechanisms from postmortem studies. J Neurochem [Internet]. 2016 Jun 15 [cited 2023 Dec 18];138 Suppl 1:54–70. Available from: https://europepmc.org/article/med/27306735

9. Kawakami I, Arai T, Hasegawa M. The basis of clinicopathological heterogeneity in TDP-43 proteinopathy. Acta Neuropathol [Internet]. 2019;138(5):751–70. Available from: https://www.ncbi.nlm.nih.gov/pubmed/31555895

10. Hasegawa M, Arai T, Nonaka T, Kametani F, Yoshida M, Hashizume Y, et al. Phosphorylated TDP-43 in frontotemporal lobar degeneration and amyotrophic lateral sclerosis. Ann Neurol [Internet]. 2008;64(1):60–70. Available from: https://www.ncbi.nlm.nih.gov/pubmed/18546284

11. Wils H, Kleinberger G, Janssens J, Pereson S, Joris G, Cuijt I, et al. TDP-43 transgenic mice develop spastic paralysis and neuronal inclusions characteristic of ALS and frontotemporal lobar degeneration. Proc Natl Acad Sci U S A [Internet]. 2010;107(8):3858–63. Available from: https://www.ncbi.nlm.nih.gov/pubmed/20133711

12. Becker LA, Huang B, Bieri G, Ma R, Knowles DA, Jafar-Nejad P, et al. Therapeutic reduction of ataxin-2 extends lifespan and reduces pathology in TDP-43 mice. Nature [Internet]. 2017;544(7650):367–71. Available from: https://www.ncbi.nlm.nih.gov/pubmed/28405022

13. Iñesta Vaquera F, Ferro F, McMahon M, Henderson CJ, Wolf CR. Potential of in vivo stress reporter models to reduce animal use and provide mechanistic insights in toxicity studies. F1000Res [Internet]. 2022 [cited 2023 Oct 24];11. Available from: https://pubmed.ncbi.nlm.nih.gov/37427015/

14. McMahon M, Frangova TG, Henderson CJ, Wolf CR. Olaparib, Monotherapy or with Ionizing Radiation, Exacerbates DNA Damage in Normal Tissues: Insights from a New p21 Reporter Mouse. Mol Cancer Res [Internet]. 2016;14(12):1195–203. Available from: https://www.ncbi.nlm.nih.gov/pubmed/27604276

15. McMahon M, Ding S, Acosta-Jimenez LP, Frangova TG, Henderson CJ, Wolf CR. Measuring in vivo responses to endogenous and exogenous oxidative stress using a novel haem oxygenase 1 reporter mouse. J Physiol [Internet]. 2018;596(1):105–27. Available from: https://www.ncbi.nlm.nih.gov/pubmed/29086419

16. Inesta-Vaquera F, Navasumrit P, Henderson CJ, Frangova TG, Honda T, Dinkova-Kostova AT, et al. Application of the in vivo oxidative stress reporter Hmox1 as mechanistic biomarker of arsenic toxicity. Environ Pollut [Internet]. 2020/11/21. 2021;270:116053. Available from: https://www.ncbi.nlm.nih.gov/pubmed/33213951

17. McMahon M, Ding S, Acosta-Jimenez LP, Frangova TG, Henderson CJ, Wolf CR. Measuring in vivo responses to endogenous and exogenous oxidative stress using a novel haem oxygenase 1 reporter mouse. J Physiol [Internet]. 2018;596(1):105–27. Available from: https://www.ncbi.nlm.nih.gov/pubmed/29086419

18. McCloy RA, Rogers S, Caldon CE, Lorca T, Castro A, Burgess A. Partial inhibition of Cdk1 in G 2 phase overrides the SAC and decouples mitotic events. Cell Cycle [Internet]. 2014 May 1 [cited 2023 Nov 6];13(9):1400–12. Available from: https://pubmed.ncbi.nlm.nih.gov/24626186/

19. Pickles S, Gendron TF, Koike Y, Yue M, Song Y, Kachergus JM, et al. Evidence of cerebellar TDP-43 loss of function in FTLD-TDP. Acta Neuropathol Commun [Internet]. 2022 Dec 1 [cited 2023 Nov 6];10(1). Available from: https://pubmed.ncbi.nlm.nih.gov/35879741/

20. Chipika R, Mulkerrin G, Pradat PF, Murad A, Ango F, Raoul C, et al. Cerebellar pathology in motor neuron disease: neuroplasticity and neurodegeneration. Neural Regen Res [Internet]. 2022 Nov 1 [cited 2023 Nov 29];17(11):2335. Available from: /pmc/articles/PMC9120698/

21. Kabiljo R, Iacoangeli A, Al-Chalabi A, Rosenzweig I. Amyotrophic lateral sclerosis and cerebellum. Scientific Reports 2022 12:1 [Internet]. 2022 Jul 22 [cited 2023 Nov 29];12(1):1–4. Available from: https://www.nature.com/articles/s41598-022-16772-5

22. Chaytow H, Carroll E, Gordon D, Huang YT, van der Hoorn D, Smith HL, et al. Targeting phosphoglycerate kinase 1 with terazosin improves motor neuron phenotypes in multiple models of amyotrophic lateral sclerosis. EBioMedicine [Internet]. 2022 Sep 1 [cited 2023 Oct 24];83. Available from: https://pubmed.ncbi.nlm.nih.gov/35963713/

23. Lin Z, Kim E, Ahmed M, Han G, Simmons C, Redhead Y, et al. MRI-guided histology of TDP-43 knock-in mice implicates parvalbumin interneuron loss, impaired neurogenesis and aberrant neurodevelopment in amyotrophic lateral sclerosis-frontotemporal dementia. Brain Commun [Internet]. 2021 Apr 5 [cited 2023 Dec 11];3(2). Available from: 10.1093/braincomms/fcab114

24. Baradaran-Heravi Y, Van Broeckhoven C, van der Zee J. Stress granule mediated protein aggregation and underlying gene defects in the FTD-ALS spectrum. Neurobiol Dis [Internet]. 2019;134:104639. Available from: https://www.ncbi.nlm.nih.gov/pubmed/31626953

25. McCauley ME, Baloh RH. Inflammation in ALS/FTD pathogenesis. Acta Neuropathol [Internet]. 2019;137(5):715–30. Available from: https://www.ncbi.nlm.nih.gov/pubmed/30465257

26. Escher BI, Henneberger L, König M, Schlichting R, Fischer FC. Cytotoxicity Burst? Differentiating Specific from Nonspecific Effects in Tox21 in Vitro Reporter Gene Assays. Environ Health Perspect [Internet]. 2020 Jul 1 [cited 2023 Oct 24];128(7):1–10. Available from: https://pubmed.ncbi.nlm.nih.gov/32700975/

27. Luan W, Wright AL, Brown-Wright H, Le S, San Gil R, Madrid San Martin L, et al. Early activation of cellular stress and death pathways caused by cytoplasmic TDP-43 in the rNLS8 mouse model of ALS and FTD. Molecular Psychiatry 2023 [Internet]. 2023 Apr 3 [cited 2023 Oct 24];1–17. Available from: https://www.nature.com/articles/s41380-023-02036-9

28. de Mena L, Lopez-Scarim J, Rincon-Limas DE. TDP-43 and ER Stress in Neurodegeneration: Friends or Foes? Front Mol Neurosci [Internet]. 2021 Oct 25 [cited 2023 Oct 24];14. Available from: https://pubmed.ncbi.nlm.nih.gov/34759799/

29. Tarot P, Lasbleiz C, Liévens JC. NRF2 signaling cascade in amyotrophic lateral sclerosis: bridging the gap between promise and reality. Neural Regen Res [Internet]. 2024 May 1 [cited 2023 Oct 24];19(5):1006–12. Available from: https://pubmed.ncbi.nlm.nih.gov/37862202/

